# capC-MAP : A Software Package for Analysis of Capture-C data

**DOI:** 10.1101/456160

**Authors:** Adam Buckle, Nick Gilbert, Davide Marenduzzo, Chris A. Brackley

**Affiliations:** SUPA, School of Physics and Astronomy, University of Edinburgh, Peter Guthrie Tait Road, Edinburgh EH9 3FD, United Kingdom; Medical Research Council Human Genetics Unit, Medical Research Council Institute of Genetics & Molecular Medicine, University of Edinburgh, Western General Hospital, Edinburgh, United Kingdom

## Abstract

We present capC-MAP, a software package for the analysis of Capture-C data. Capture-C is a “many-to-all” chromosome-conformation-capture method. We summarise the method, then detail capC-MAP, the first software specifically designed and optimised for Capture-C data. capC-MAP has been developed with ease-of-use and flexibility in mind: the entire pipe-line can be run with a single command, or the component programs can be run individually for custom data processing, in a strategy that will suit computational as well as experimental researchers. Finally, we compare and benchmark capC-MAP against another package which can perform (though is not optimised for) analysis of Capture-C data.

## Introduction

Over recent years the family of experimental methods based on chromosome-conformation-capture (3C) has grown [1], with different variants used to generate data at different resolutions, in populations and single cells, and using different methods of detection – e.g. PCR, microarray, or next-generation sequencing (NGS) technologies. These methods are used to probe the interactions between different chromatin regions *in vivo,* and to uncover the three-dimensional organization of chromosomes and genomes. In the near two decades since they were first developed, they have revolutionised our understanding of genome organisation and functions [2].

The 3C based protocols range from the original “one-to-one” 3C method [3] which measures interactions between selected pairs of genomic loci, through the “one-to-all” style 4C method [4], where genome wide interactions for a single selected loci are obtained, to high-throughput “all-to-all” HiC [5], which uses NGS to obtain genome-wide chromatin interaction maps. Capture-C is a relatively recent addition to the 3C family, developed by Hughes *et al.* and first reported in Ref. [6]; it uses oligo-capture technologies, a frequently cutting restriction enzyme, and NGS sequencing, to deliver high-resolution cis-interaction profiles for up to hundreds of target loci from a single experiment. While HiC can provide a large-scale overview of chromosome interactions, deep sequencing is required to get good spatial resolution, which is costly. Capture-C is a “many-to-all” assay which gives interaction profiles for a set of “targets” at near restriction enzyme fragment resolution [6, 7]. The targeted approach means that high-resolution data is achievable with a fraction of the amount of sequencing than is required for HiC, and with a smaller amount of starting material, making the method applicable to small samples and difficult to obtain tissues [7].

Popularity of the Capture-C method has grown [8–17], but the analysis of the data is complicated – it requires non-standard use of bioinformatics tools as well as some bespoke data treatment. Most work using the method to date has used custom analysis scripts accessible only to experts in bioinformatics and programming. While some analysis tools designed to treat HiC data now also support Capture-C, these are not optimized for the method and are limited in functionality, and there has been a lack of easy to use, dedicated software tools. Here we introduce capC-MAP, a software package for the analysis of Capture-C (or NG Capture-C [7]) data. This is the first package dedicated to Capture-C experiments, and has been designed with both ease of use and flexibility in mind. The software comprises a suit of programs written in Python and C**++**, with a Python wrapper script which enables a whole Capture-C experiment analysis to be run with a single command. capC-MAP automates all of the processing steps, including calling external standard bioinformatics tools. It is designed to be easy to use by anyone familiar with Unix-like operating systems, but the individual component programs can also be used independently by more experienced users in custom pipe-lines.

In this paper we first describe the Capture-C method, we then detail capC-MAP and the different processing steps it performs, before comparing our software with other (non-dedicated) tools which can handle Capture-C data.

## Results

### The Capture-C method

The Capture-C protocol is described in Refs. [6, 7]; for completeness we summarize the main details of a typical experiment and introduce some terminology.

The underlying principles of all 3C based methods are similar and summarised in Fig. 1a. First, formaldehyde fixation is used to cross-link proteins and DNA within intact nuclei, physically linking DNA segments which are in close spatial proximity. Next the formaldehyde cross-linked DNA template is digested in the intact nuclei [18] with a selected restriction enzyme, and then the DNA is re-ligated. Since ligation is likely to occur between cross-linked fragments, this results in the joining of fragments that were not adjacent in the linear genome, but were close together in 3-D space. The resulting DNA is purified to form a 3C library. In the Capture-C method the 3C library is then sonicated to an optimal size of ~ 300 bp, and used to prepare sequencing libraries, whereupon solution-based sequence capture technology is used to enrich for certain restriction enzyme fragments. Specifically, biotin labelled RNA or DNA capture oligos are designed against a set of restriction fragments of interest – these hybridise with the DNA fragments, which are then pulled down with the biotin tag, before re-amplification using primers to sequencing adapters. Since the library consists of hybrid fragments representing the proximity ligation events, paired-end sequencing reveals which distal fragments were in proximity to the fragments of interest (Fig. 1b). Here we use the terminology “targets” to refer to the restriction enzyme fragments for which oligos have been designed, and “reporters” to refer to any fragments which have been found ligated to a target. The set of reporters for a given target can be used to build up a picture of the interactions genome wide (they are “piled-up” to provide an “interaction profile”). The data generated from Capture-C is similar to 4C, but here the capture oligos provide the viewpoints, so multiple interaction profiles can be obtained from a single experiment (note that our term *target* is synonymous with the *viewpoint* or *bait* in 4C).

**Fig. 1:**
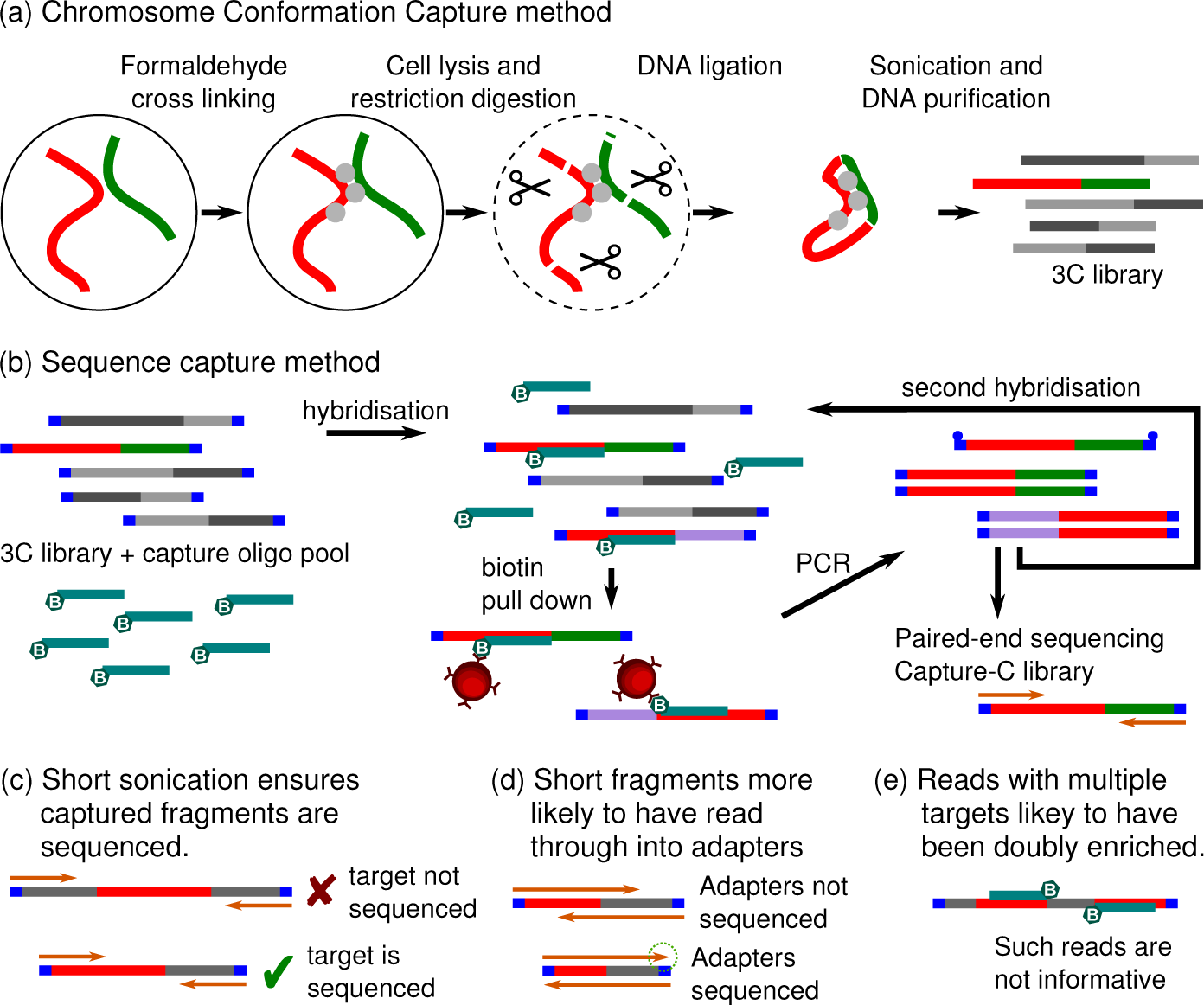
Schematic showing the steps of the Capture-C method. (**a)** 3C library preparation steps: formaldehyde cross-linking; cell lysis; digestion of cross-linked chromatin by the *Dpn*II restriction endonuclease; ligation by T4 DNA ligase, joining hybrid DNA fragments together; and finally, DNA sonication and purification to produce a 3C library. (**b)** Sequence capture steps of the Capture-C methodology: Illumina sequencing adapters are added (blue); an experiment specific biotin labelled capture oligo pool is combined with the 3C library in a hybridisation reaction; streptavidin beads (red circles) are used to pull down biotin oligo/3C DNA complexes; and then the library is re-amplified using primers to the Illumina sequencing adapters. After this, a second round of hybridisation and amplification can be performed before paired-end sequencing. For further details see Refs. [6, 7]. (**c-e)** Some technical points of the Capture-C method. Sonication to a small size (200-300 bp, similar to the length of *Dpn*II restriction fragments) is recommended to ensure that the captured targets are sequenced. Paired-end sequencing of short fragments is likely to lead to adapter contamination as a result of read through into 3′ end. Ligation fragments with multiple targets may have been captured multiply; since oligo efficiency is generally unknown, such fragments are not quantitatively informative.

Capture-C uses a restriction endonuclease with a four base-pair recognition sequence (typically *Dpn*II); the short recognition sequence means it appears frequently within the genome, resulting in short restriction fragments. This – together with the oligo hybridisation step which vastly improves the signal to noise ratio by reducing the number of background ligation events being sequenced – leads to very high-resolution data. The short restriction fragment size necessitates that the library be sonicated to a similar short length (compared to that typical of HiC experiments) to ensure that the captured fragment falls within the sequenced region (see Fig. 1c).

There are several possible oligo design strategies, but typically this entails designing oligos which bind each end of the target fragments. In the original Capture-C method [6] Hughes *et al.* used RNA oligos synthesized on a microarray (meaning that the design included a minimum of 40,000 oligos), designed such that each end of each target fragment was tiled by several oligos. In a modified version of the method (named NG Capture-C [7]) a single 120 bp biotinylated DNA oligo was designed for each end of each target fragment – this allowed for a more cost effective and scalable experimental design. The Hughes lab developed an on-line oligo design tool called CapSequm (http://apps.molbiol.ox.ac.uk/CaptureC/cgibin/CapSequm.cgi) which performs a BLAT search [19] and Repeatmasker analysis [20] to generate robust capture oligos which will hybridise to a single restriction fragment. Another improvement in the NG Capture-C protocol is that two successive rounds of sequence capture are performed, the first giving a 5-20,000 fold enrichment, with the second able to achieve up 1,000,000-fold enrichment. This dramatically improves the signal-to-noise ratio and reduces the required sequencing depth. This efficiency also offers the ability to pool multiple 3C libraries with indexed sequencing adapters, which can then be processed in a single reaction [7].

### capC-MAP overview

In developing capC-MAP our aim was to automate the analysis of Capture-C data, going from fastq files of sequenced reads to a set of outputs for each target using a single command line. The main output from the software is an “interaction profile” for each target showing intrachromosomal interactions (interchromosomal interactions are output separately). Using an easily customisable “configuration” file the user can specify different normalization and binning options. Interaction profiles are output in the standard bedGraph format – there are many tools available for visualization and downstream analysis of data in this format. For example, IGV [21] or the UCSC genome browser [22] can be used for visualization, and the BEDtools suit [23] or many of the R packages available via bioconductor [24] can be used for downstream analysis and plotting (for example the “peakC” package performs non-parametric peak calling on 4C and Capture-C data [25]).

### capC-MAP details

The analysis pipe-line which capC-MAP follows is based on that detailed in Ref. [6], and is shown schematically in Fig. 2. Here we summarise these steps.

- Since it is recommended that during library preparation fragments are sonicated to an average length of 200-300 bp, it is likely that read through into the adapter sequence will have occurred during paired-end sequencing (see Fig. 1d). The first step in the analysis is therefore to trim adapter sequence from the mapped reads – this is done using the **cutadapt** software [27] (trimming the common adapter sequence).
- Next we perform an *in silico* restriction enzyme digestion; i.e. each read-pair is searched for instances of the enzyme cut sequence (GATC for *Dpn*II), and is broken into smaller fragments at these positions. Thus a group of read fragments is obtained from the read pair.
- The group of fragments is then aligned to the reference genome using the **bowtie** software [26], as though they were single-end reads (bowtie would make incorrect assumptions about valid alignments if run in paired-end mode). Since the fragments may be quite short, bowtie is run with quite stringent reporting criteria to ensure only uniquely mapped fragments are reported. This is the most time consuming step of the analysis, and can be run in parallel on a multi-core computer.
- The output from bowtie is a SAM format file containing details of all mapped (and unmapped) fragments. This needs to be sorted by read name so that the original fragment groups can be recovered. At this point duplicates are removed: these are defined as read groups where identical fragments appear in the same order, and two fragments are said to be identical if either they mapped to the same position in the genome, or they did not map but have identical sequence. Such duplicates are likely to have arisen from PCR artefacts.
- Next, the set of mapped fragments within each read group is compared to a genome wide map of restriction enzyme fragments and to the list of target fragments, in order to identify “targets” and “reporters”.
- At this point invalid interactions are identified and removed, and the remaining valid intra- and interchromosomal interactions are stored separately for each target. Invalid interactions are
  – interactions between targets; since these will have resulted from ligation events which bring together regions which bind more than one oligo they are likely to have been doubly enriched and are therefore not quantitative (Fig. 1e).
  – interactions with an “exclusion zone” around another target; since digestion is not 100% efficient, these could also have resulted from ligation events which bring together regions which bind more than one oligo.
  – interactions with multiple (non-adjacent) reporters; it is possible to find fragments which map to more than two distal chromatin regions within the same read group – though these are in theory informative [28, 29], in practice they occur very rarely, so for simplicity we treat them as invalid.
- Finally, for each target the list of intrachromosomal interactions is “piled-up” to generate a bedGraph format file which counts the number of interactions between that target and every other restriction enzyme fragment in the same chromosome. This gives a raw restriction enzyme fragment level interaction profile. Depending on the read coverage it can also be useful to generate a binned and smoothed interaction profiles (detailed below).

**Fig. 2:**
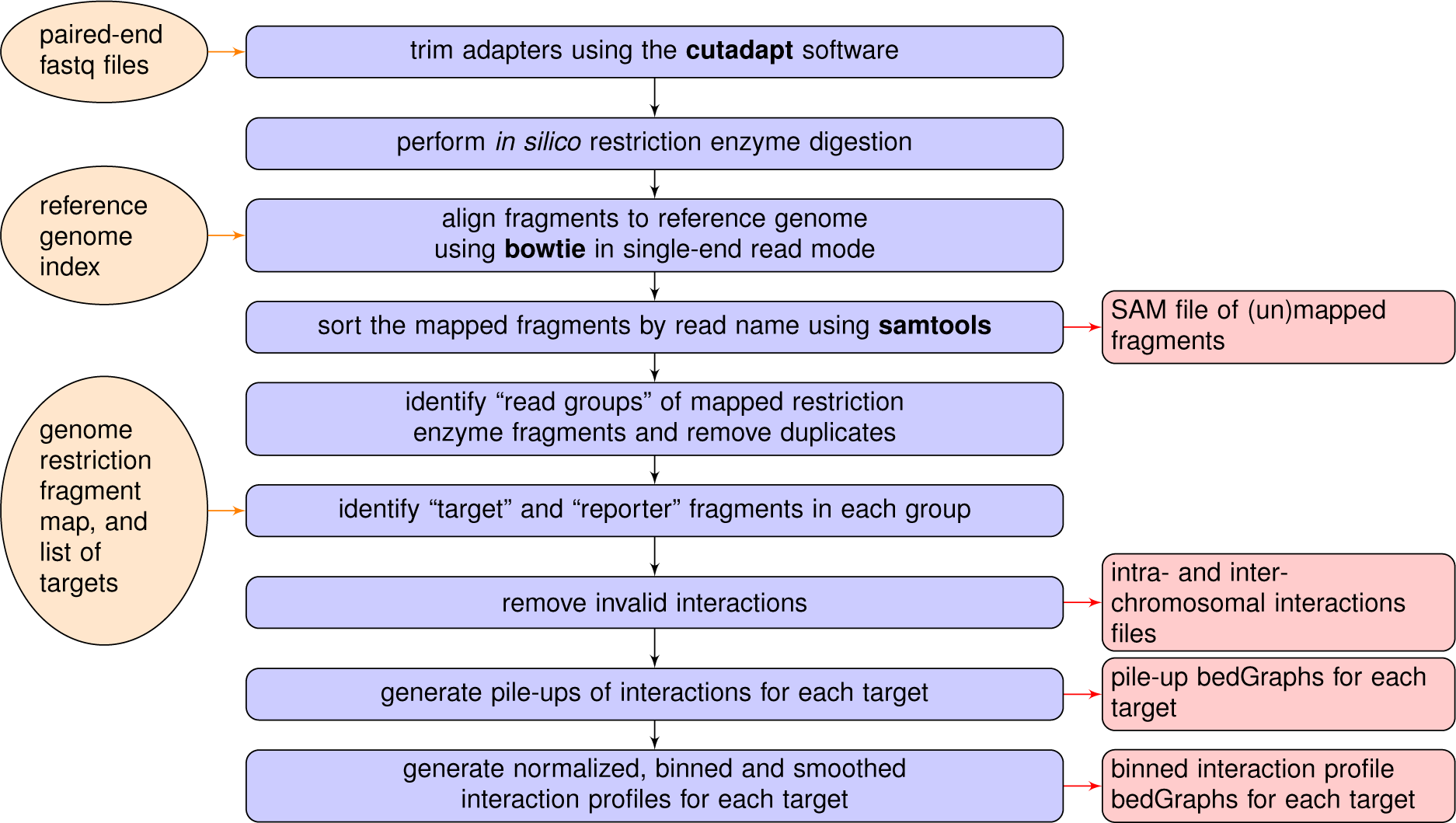
Flow diagram for the Capture-C analysis process. Each of the steps completed by the capC-MAP software during a typical run is shown (centre). Where external software is called this is shown in bold. Required inputs are shown to the left (the reference genome index is generated using the bowtie alignment software [26], and the genome-wide map of restriction enzyme fragments can be generated by capC-MAP). Typical outputs are shown to the right.

capC-MAP outputs a set of informative log files and intermediate data files as detailed in the documentation, but the two main types of output for each target are raw pile-ups of interactions, and interactions profiles which have been binned and smoothed. These are both in the standard bedGraph file format. The raw pile-up files contain a line for each restriction fragment with which a target interacts, with a count for the number of times this interaction was observed. A typical plot of this data is shown in Fig. 3a. capC-MAP can also provide normalized output in units of “reads per million”, i.e. the read counts are normalized such that for each target they sum to one million genome wide.

**Fig. 3:**
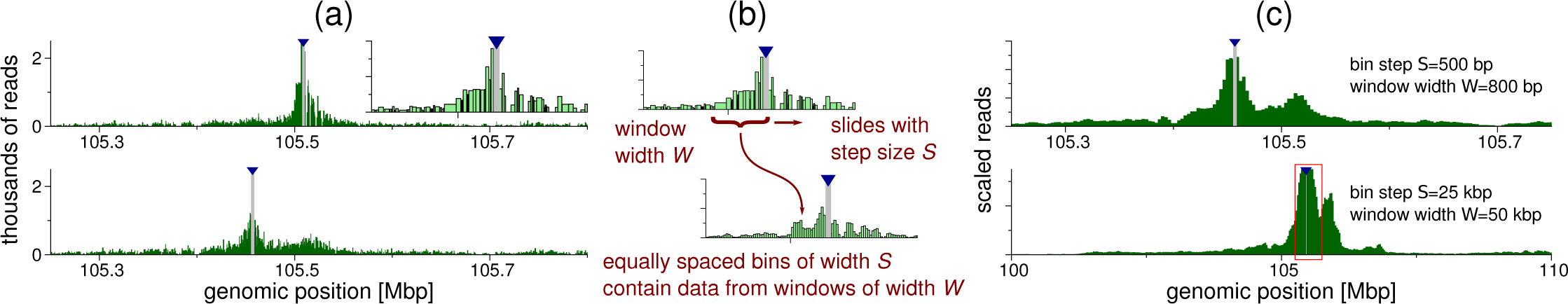
Typical interaction profiles from a Capture-C experiment. (**a)** Plots showing raw pile-ups of reads for two different targets from experimental data published in Ref. [17] (these are targets on chromosome 2 of mouse mm9 build; data obtained from GEO:GSE120666). The grey bar and blue arrowhead indicate the position of the target. Inset shows a zoom around the target shown in the top plot – here it is evident that restriction fragments are of different sizes. (**b)** Schematic showing the sliding window binning scheme used by capC-MAP. (**c)** Plots showing binned smoothed profiles generated from the data shown in the bottom plot of panel (a), for two different choices for the window width and bin step size. The red box in the bottom plot shows the region displayed in the top. Note that capC-MAP does not provide functionality to generate plots since many software tools which can read and plot bedGraph files are available.

It is often useful to apply some binning or smoothing to interaction profiles, and capC-MAP provides options to do this. For example, if a data set has a low read count per target, binning can give smoother and easier to interpret interaction profiles. Also, since the length of restriction fragments has quite a broad distribution, examining raw pile-ups can be misleading; e.g, if we consider two regions which interact with a target at a similar frequency – if one region has a single long restriction enzyme fragment, and the other has several short ones, in the latter case the same number of interactions would be shared across more fragments, resulting in a lower ‘per fragment’ interaction count. capC-MAP uses sliding window binning, where the user specifies a window width *W* and a step size *S,* where *W* ≥ *S.* Bins go up in steps of size *S* bp, and each contains the number of reads within a window of width *W* bp around the bin centre (this strategy is shown schematically in Fig.3b). It is often useful to generate profiles with several different bin/window size combinations, e.g. depending on whether short or longer ranged interactions are of interest; Fig.3c shows examples of different binned and smoothed interaction profiles.

### Comparison with other software

To our knowledge, the only other publicly available software which automates analysis of Capture-C data is HiC-Pro [30]. This is a popular tool for HiC data analysis, and a recent update added the ability to analyse Capture-C data. In order to compare the efficiency of HiC-Pro and capC-MAP we used a data set from Ref. [7] as a test case, running an analysis on the same machine with each software package. In both cases two processor cores were used, and all standard options selected (HiC-Pro was run in ‘sequential mode’ where some processing steps which are only relevant for HiC data were skipped). Some details of the analysis from each software package are shown in Table 1; note that the packages may count reads in different ways meaning that not all quantities are directly comparable. We find that capC-MAP identifies a higher proportion of PCR duplicates, and finds approximately 1.7 times more informative reads; also capC-MAP performs the analysis in under a third the time taken by HiC-Pro, highlighting that the packages are optimized for different types of data. A major difference between the two pipelines is that capC-MAP performs an *in silico* digestion of the reads before alignment of the resulting fragments, whereas HiC-Pro attempts alignment before digestion and only searches for enzyme cut sites if this fails (i.e., where a ligation junction appears within the read, alignment is attempted multiple times). The latter strategy is optimal for HiC data where typically a less frequently cutting enzyme is used (e.g. *Hind*III) and the ligation fragments tend to be longer, meaning the sequenced regions are less likely to contain a cut site; in the case of Capture-C using *Dpn*II, the fragments are highly likely to include a cut site, so for most fragments HiC-Pro will need to attempt alignment multiple times. Another possible reason for the lower efficiency of HiC-Pro is that it uses the Bowtie2 aligner [31], whereas capC-MAP uses Bowtie1 which is optimised for short reads [26]. We also note that while HiC-Pro provides a utility to extract interaction profiles from the data set, there is no functionality for binning or smoothing the data.

**Table 1:** Table comparing output from capC-MAP v0.0.1 and HiC-Pro v2.11 [30]. A data set from Ref. [7] (GEO:GSE67959) was used as a test case, run using two cores of a 10 core hyper-threaded 2.6GHz Intel Xeon E5-2660 processor machine with 50GB RAM running Scientific Linux 7.5. Note that the two software packages report read statistics in different ways, so may not be directly comparable, and not all values are available from both.

| Data set details |  |
| --- | --- |
| Total reads | 37,381,686 |
| Number of target fragments | 35 |

**Table 1: Table comparing output from capC-MAP v0.0.1 and HiC-Pro v2.11 [30].**
|  | capC-MAP | HiC-Pro |
| --- | --- | --- |
| number of duplicates removed | 3,391,919 (9.07 %) | 485,560 (1.29 %) |
| number of reads without a mapped target-reporter pair | 32,659,794 | – |
| number of invalid interactions removed | 172,596 | – |
| total informative reads | 1,140,683 | 667,259 |
| of which were interchromosomal | 290,516 | 106,348 |
| of which were intrachromosomal | 850,167 | 560,911 |
| average interchromosomal per target | 8,300 | 3,038 |
| average intrachromosomal per target | 24,290 | 16,026 |
| total run time | 3 hours 31 mins | 11 hours 39 mins |

capC-MAP was designed with the intention of extracting interaction profiles (such as might be obtained in a 4C experiment) for each targeted restriction enzyme fragment. Other experimental designs include (i) capture oligos designed to tile a region or chromosome of interest to obtain a HiC-style map (as in Ref. [32]); and (ii) oligos designed to capture interactions from many thousands of dispersed sites (e.g. all promoters), to identify significant interactions between target and non-target or between pairs of target cites (as in Refs. [33, 34]). These approaches often use the “Capture Hi-C” protocol [33, 35, 36] which combines elements from the Capture-C and Hi-C methods. For such experiments, analysis strategies different from those employed by capC-MAP might be more relevant, and tools such as HiC-Pro [30] and CHiCAGO [36] (which is designed specifically for experiments of type (ii)) may be appropriate. We note that in design case (ii) it is possible to use capC-MAP to generate interaction profiles for each target, though their quality will depend on the read depth (which may be lower than a Capture-C experiment if the same number of reads are diluted across thousands of targets).

## Conclusions

In this paper we present capC-MAP, a software package for the analysis of Capture-C data. The method, which yields “many-to-all” type chromosome conformation capture information, has become increasingly popular [8–17], but until now has been restricted by a lack of an easy-to-use data analysis software package. capC-MAP allows the entire processing pipeline to be performed using a single command line, but also provides a suite of tools for advanced and bespoke analysis. Here we have compared the function of capC-MAP with another similar package, showing that the Capture-C specific optimizations present in capC-MAP give rise to a better yield of informative interactions and a three-fold shorter processing time.

### Materials and Methods

capC-MAP is implemented as a suit of programs written in **C++** and Python (version 2.7). Each program can be run separately, or an entire analysis can be performed with a single command line. capC-MAP also calls the external software packages cutadapt [27], bowtie [26] and samtools [37], which are freely available open-source software. capC-MAP should run on any standard Unix-style system (including Linux and Mac) where the above listed software is installed.

For a typical Capture-C experiment, the user will need to perform the following steps:

1. Build an index for the reference genome for the bowtie alignment software;
2. Build a restriction enzyme fragment map for the reference genome using the capC-MAP “genomedigest” tool;
3. Run the full analysis pipe line using the capC-MAP “run” tool;

where steps 1 and 2 only need to be performed once for each reference genome. Step 3 requires only a single command line, and the software reads a “configuration file” to set all the options. A template configuration file is provided with the software, and this is straightforward to modify for a specific experiment. capC-MAP can run some steps in parallel by taking advantage of the shared memory multi-threading options of the external programs it calls (since the slowest steps of the analysis are aligning the reads to the reference genome, and sorting the resulting SAM file, only these can be performed in parallel). capC-MAP also provides functionality for handling targets which appear at multiple points in a genome, and for combining replicate experiments.

Full usage details are provided in the documentation which accompanies the software. capC-MAP is available under the GNU General Public License v3.0, and its source code is available at https://github.com/cbrackley/capC-MAP. Full documentation is available at capc-map.readthedocs.io. The data used to generate Fig. 3 was obtained from GEO:GSE120666. The data set used to perform the benchmarking detailed in Table 1 was obtained from GEO: GS E67959.

## Acknowledgements

The work was supported by the European Research Council for funding (Consolidator Grant THREEDCELLPHYSICS, Ref. 648050), and research in the Gilbert lab is funded by the UK Medical Research Council (MR/J00913X/1)

